# HSFs drive stress type-specific transcription of genes and enhancers

**DOI:** 10.1101/2021.11.23.469701

**Authors:** Samu V Himanen, Mikael C Puustinen, Alejandro J Da Silva, Anniina Vihervaara, Lea Sistonen

## Abstract

Reprogramming of transcription is critical for the survival under cellular stress. Heat shock has provided an excellent model to investigate nascent transcription in stressed cells, but the molecular mechanisms orchestrating RNA synthesis during other types of stress are unknown. We utilized PRO-seq and ChIP-seq to study how Heat Shock Factors, HSF1 and HSF2, coordinate transcription at genes and enhancers upon oxidative stress and heat shock. We show that pause-release of RNA polymerase II (Pol II) is a universal mechanism regulating gene transcription in stressed cells, while enhancers are activated at the level of Pol II recruitment. Moreover, besides functioning as conventional promoter-binding transcription factors, HSF1 and HSF2 bind to stress-induced enhancers to trigger Pol II pause-release from poised gene promoters. Importantly, HSFs act at distinct genes and enhancers in a stress type-specific manner. HSF1 binds to many chaperone genes upon oxidative and heat stress but activates them only in heat-shocked cells. Under oxidative stress, HSF1 and HSF2 *trans*-activate genes independently of each other, demonstrating, for the first time, that HSF2 is a *bona fide* transcription factor. Taken together, we show that HSFs function as multi-stress-responsive factors that activate specific genes and enhancers when encountering changes in temperature and redox state.

## INTRODUCTION

Cells are exposed to various cytotoxic stresses including elevated temperatures and oxidative stress. While increased temperatures lead to protein misfolding, oxidative stress is caused by elevated production of reactive oxygen species (ROS) that oxidize macromolecules (proteins, lipids and nucleic acids) (1, 2). Regulation of ROS levels is critical for cell survival and also for normal physiology, since basal levels of ROS activate cellular signalling pathways, while increased production of ROS promotes aging and progression of many diseases, such as cancer (1, 3). To combat cytotoxic stresses, cells extensively reprogram their transcription (4). Although genome-wide transcription is repressed upon stress, certain stress-responsive transcription factors can *trans*-activate pro-survival genes, allowing cells to overcome the adverse conditions (4-6). Transcription under oxidative stress is known to be regulated by nuclear factor erythroid 2-related factor 2 (Nrf2) and forkhead box transcription factors (FOXOs), while proteotoxic stress-inducible transcription is driven by a family of heat shock factors (HSFs) (4). In addition to gene activation, cytotoxic conditions have been shown to activate transcription at numerous enhancers, which are distal regulatory elements in the DNA that can promote gene expression through loop formation (6-10). Intriguingly, active enhancers produce short and unstable enhancer RNAs (eRNAs) that regulate gene transcription by mechanisms which are not entirely understood (10). The characteristic pattern of eRNA transcription serves as a means to identify active enhancers de novo using methods that measure nascent transcription at a nucleotide resolution (11-13).

The master *trans*-activators in stressed cells include the HSFs, which are activated in response to various proteotoxic stresses, e.g. heat shock (14, 15). Proteotoxic stress impairs proper protein folding and causes accumulation of unfolded proteins (2). To prevent and mitigate these damages, HSFs rapidly *trans*-activate genes encoding heat shock proteins (HSPs), which, in turn, function as molecular chaperones (4). HSF1 is the master regulator of chaperone expression and the most studied member of the HSF family, whereas HSF2 has been mainly characterized as a developmental transcription factor, particularly in gametogenesis and neurogenesis (15). Intriguingly, exogenous human HSF2, but not HSF1, can substitute for yeast HSF to provide thermotolerance, demonstrating that HSF2 has a capability to act as a stress-responsive transcription factor (16). There is also evidence for a context-dependent interplay between HSF1 and HSF2, either competitive or synergistic, but the functional role of HSF2 in stress-inducible transcription has remained elusive (17). Although HSF1 has been identified as the master regulator of the heat shock response and other proteotoxic stresses, it is also activated in response to oxidative stress (18). The biological significance of HSF1 in the regulation of redox status was previously reported in a study, where increased production of cardiac ROS was observed in the absence of HSF1 (19). Nevertheless, how HSF1 and other member of the HSF family contribute to transcriptional reprogramming upon oxidative stress is unknown.

Recently, it was shown that apart from binding promoters, HSF1 is recruited to heat-induced enhancers to activate genes, such as forkhead box O3 (*Foxo3*) and tax1-binding protein 1 (*Tax1bp1*) (6, 9, 20). The function of the HSF family members in the genome-wide enhancer activation under different stress conditions is, however, not known. In this study, we compared the stress-specific transcription programs by tracking transcription at genes and enhancers in cells exposed to either oxidative stress or heat shock. We used precision run-on sequencing (PRO-seq), which quantifies transcriptionally engaged RNA polymerase II (Pol II) complexes at a single nucleotide resolution across the genome (11). Unlike RNA-seq and other conventional methods that measure steady-state mRNA levels, PRO-seq allows detection of active transcription at promoter-proximal regions, upstream divergent transcripts, gene bodies, termination windows, and enhancers (11, 12, 21). Combining PRO-seq with chromatin immunoprecipitation sequencing (ChIP-seq), we identified HSF1 and HSF2 as new regulators of oxidative stress-inducible transcription. HSF1 and HSF2 were recruited to distinct genomic sites in cells exposed to oxidative stress or heat shock, which triggered the activation of stress-specific transcription programs. Previously, HSF2 has been characterized as a modulator of HSF1 activity upon heat shock, but here we found that during oxidative stress, HSF2 regulates transcription independently of HSF1. Furthermore, besides functioning as conventional promoter-binding transcription factors, both HSFs were required for the activation of several oxidative stress- and heat-inducible enhancers. Finally, we found that in contrast to the promoter-bound HSF1, which drives the classical chaperone genes, binding of HSF1 to enhancers activates genes encoding proteins localized at plasma membrane and cell junctions. Taken together, our results show that HSFs function as multi-stress-responsive transcription factors that orchestrate stress-specific transcription programs through genes and enhancers.

## MATERIAL AND METHODS

### Cell lines

Wild-type (WT) and HSF1 knock-out (KO) MEFs were derived from mice generated in the laboratory of Ivor Benjamin (45). HFS2 KO MEFs were from mice generated in the laboratory of Valerie Mezger (46).

### Cell culture and treatments

MEFs were grown in Dulbecco’s modified eagle medium (Sigma) supplemented with 10 % fetal bovine serum, 2 mM L-glutamine, 50 μg/ml penicillin/streptomycin, and non-essential amino acids (Gibco). Cells were maintained at 37°C with 5 % CO_2_. Cells were exposed to heat shock by submerging the cell culture dishes into a 42°C water bath for an hour. Oxidative stress was induced by treating the cells with 30 μM freshly prepared menadione solution at 37°C. DNA damage was induced by exposing cells to 2 mM hydroxyurea for 17 h.

### Western Blotting

Cells were lysed in laemmli sample buffer (30 % glycerol; 3 % SDS; 188 mM Tris-Cl, pH 6.8; 0.015 % bromophenol blue; 3 % β-mercaptoethanol). Equal volumes of lysates were run on SDS-PAGE, after which proteins were transferred to nitrocellulose membrane. Membranes were blocked with nonfat dried milk diluted in PBS-Tween20 for 1 hour at room temperature (RT). Proteins bound to membrane were analyzed using primary antibodies against HSF1 (ADI-SPA-901, Enzo), HSF2 (3E2, EMD Millipore) and β-tubulin (T8328, Merck). Next, the membranes were incubated in secondary HRP-conjugated antibodies, and the proteins were detected with enhanced chemiluminescence.

### Immunofluorescence

WT MEFs were plated on MatTek plates (P35GC-1.5-14-C, MatTek Corporation) 48 h before treatments. Cells were fixed with 4% paraformaldehyde (PFA) for 10 min, permeabilized in 0.1% Triton X-100 in PBS and washed three times with PBS. Samples were blocked with 10% FBS in PBS for 1 h at RT and incubated overnight at 4°C with a primary anti-γH2AX antibody (05-636, EMD Millipore, 1:500 in 10% FBS-PBS). Following primary antibody incubations, the samples were washed three times with PBS. Next, samples were incubated in a secondary goat anti-mouse Alexa Fluor488 antibody (A11001, Invitrogen, 1:500 in 10% FBS-PBS) for 1 h at RT. Finally, the samples were washed two times with PBS, incubated with 300 nM DAPI diluted in PBS, and covered with VECTASHIELD mounting medium (H-1000, Vector Laboratories). All images were acquired with a 3i CSU-W1 spinning disc confocal microscope (Intelligent Imaging Innovations).

### Measurement of GSH/GSSG ratio

The effect of menadione on the induction of oxidative stress was determined by measuring the ratio between oxidized and reduced glutathione (GSH/GSSG) using a commercial kit by Promega (GSH/GSSG-Glo Assay, V6611).

### PRO-seq

PRO-seq was performed from two biological replicates as described previously (11, 47). Nuclei of MEFs were isolated in buffer A (10 mM Tris-HCl pH 7.4, 300mM sucrose, 3 mM CaCl_2_, 2 mM MgCl_2_, 0.1% Triton X-100, 0.5 mM DTT) using a dounce homogenizer. The isolated nuclei were flash-frozen and stored at −80°C in a storage buffer (10 mM Tris-HCl pH 8.0, 25% glycerol, 5 mM MgCl_2_, 0.1 mM EDTA, 5 mM DTT). Run-on reactions were performed at 37°C for 3 minutes in the presence of biotinylated nucleotides (5 mM Tris-HCl pH 8.0, 2.5 mM MgCl_2_, 150 mM KCl, 0.5 mM DTT, 0.5 % Sarkosyl, 0.4 u/μl RNase inhibitor, 0.025 mM biotin-ATP/CTP/GTP/UTP [Perkin Elmer]). Equal amounts of nuclei extracted from *Drosophila* S2 cells were used as spike-in material in run-on reactions. Total RNA was isolated with Trizol, precipitated with ethanol and fragmented by base hydrolysis using NaOH. Biotinylated transcripts were isolated with streptavidin-coated magnetic beads (M280, Invitrogen). In the next steps, TruSeq small-RNA adaptors were ligated to the ends of nascent RNAs. Before ligating 5’adaptor, the 5’-cap was removed with RNA 5’ pyrophosphohydrolase (Rpph, NEB), after which 5’end was repaired with T4 polynucleotide kinase (NEB). Nascent RNAs containing the adaptors were converted to cDNA, amplified by PCR and sequenced using NovaSeq 6000. The raw files are available in GEO accession: GSE183245.

### ChIP-seq

HSF1- and HSF2-bound DNA fragments were isolated using ChIP as previously described (32). Briefly, cells were crosslinked with 1 % paraformaldehyde for 5 min, after which paraformaldehyde was quenched with 125 mM glycine. Cells were lysed and the chromatin was fragmented by sonication with Bioruptor Pico (Diagenode) using seven cycles (30 sec on/off). Agarose gel electrophoresis was used to verify that fragment size after sonication was 300–400 bp. The following antibodies were used for immunoprecipitation: HSF1 (ADI-SPA-901, Enzo), HSF2 (Östling et al., 2007) and normal rabbit IgG (EMD Millipore). Crosslinks were reversed by incubating the samples at 65°C overnight, and the DNA was purified with phenol:chloroform. ChIP-seq libraries were generated using NEXTFLEX ChIP-seq kit and barcodes (Perkin Elmer). NovaSeq 6000 was used to sequence ChIP-seq libraries. The raw files are available in GEO accession: GSE183245.

### Mapping of PRO-seq and ChIP-seq data

Adapters were removed from the sequencing reads using cutadapt (48) and the reads were mapped to mouse genome (mm10) using Bowtie 2 (49). PRO-seq reads were mapped in single-end mode with parameters: --sensitive-local. ChIP-seq reads were mapped in paired-end mode with parameters: --sensitive-local --no-mixed --no-discordant --no-unal. The raw data (GSE183245) is available in Gene Expression Omnibus database (https://www.ncbi.nlm.nih.gov/geo/).

### Normalization of PRO-seq data

We utilized equal amounts of nuclei from *Drosophila* S2 cells as spike-in material in run-on reactions. Transcripts produced by *Drosophila* S2 nuclei are retained in the samples through every step of PRO-seq and therefore, reads mapping to *Drosophila* genome (dm3) can be used to normalize the sample data. We observed a high variance in the percentage of reads that aligned to dm3 genome between the samples. Using spike-in normalization method would have resulted in large changes to the number of upregulated and downregulated genes. Thus, we normalized the data from each sample to the total number of reads and to genes whose expression level remained the same across all samples.

### Normalization of ChIP-seq data

Spike-in normalization was utilized by adding equal amounts of chromatin from heat-shocked human Hs578T cells to each immunoprecipitation reaction. Hs578T cells were exposed to heat shock because it triggers the binding of HSF1 and HSF2 to chromatin, which in turn, allows simultaneous immunoprecipitation of HSF-bound DNA from the sample and spike-in material. We verified that each sample contained equal proportion of spike-in material by mapping the sequencing reads to human genome (hg38).

### Quantification of transcription at genes

Actively transcribed genes were identified using discriminative regulatory elements identification from global run-on data (dREG; https://dreg.dnasequence.org) (13), which detects transcription initiation sites at genes and enhancers. Genes whose TSS overlapped with dREG-called initiation sites, were retained for further analysis. Transcription was quantified from the gene bodies, which were defined as +0.5 kb from TSS to −0.5 kb from CPS. In addition, the maximum length of genes was set to 300 kb, since Pol II can only travel 240 kb during 2 hour-treatments at elongation rate of 2 kb/min (50, 51).

### Identification of transcribed enhancers

Transcribed regulatory regions, including promoters and enhancers, were identified from PRO-seq data using dREG gateway (https://dreg.dnasequence.org/). The regions that resided over 1 kb from any TSS of annotated gene, were defined as transcribed enhancers. First, we identified enhancers individually in each sample and then merged the coordinates of enhancers between the samples using bedtools merge with parameters: d −100 (52). The list of merged enhancers was used to quantify the level of enhancer transcription in individual samples.

### Differential expression analysis

Changes in transcription of genes and enhancers were determined using DESeq2 (53). Differential gene expression was measured in gene bodies, whose coordinates were defined as +0.5 kb from TSS to −0.5 kb from CPS. Changes in enhancer transcription were analyzed separately from plus and minus strands using the enhancer coordinates determined with dREG. To call statistically significant changes in transcription, p-value threshold was set to 0.05 and fold change threshold at 1.5.

### ChIP-seq peak calling

ChIP-seq peaks were identified from two combined replicates using findPeaks tools included in HOMER program (54). For HSF1 and HSF2 peaks to be called statistically significant, we set the FDR threshold to 0.001 (default value used by HOMER) and required that the fold change over IgG was at least five. For H3K27ac and H3K4me1 peaks to be statistically significant, FDR threshold was set to 0.001 and fold change over input was required to be at least four. HSF1 and HSF2 peaks were called using parameters: -style factor -F 5 -L 7 -localSize 20000. H3K27ac and H3K4me1 peaks were called using parameters: -region -L 0 -size 250.

### GO analysis

Biological processes enriched in distinct groups of HSF target genes were identified using gene ontology (GO) analysis, which is provided by GO consortium (http://geneontology.org/).

### Analysis of HSE content

Content of HSE motif in the target genes and enhancers of HSFs was analyzed using findMotifsGenome.pl tool included in HOMER program (54). HSE content was analyzed within 2 kb regions centered around the summits of HSF1 and HSF2 peaks.

### Additional datasets used

H3K27ac and H3K4me1 ChIP-seq data is from GEO dataset: GSE99009.

## RESULTS

### Oxidative stress and heat shock reprogram transcription of distinct genes and enhancers

To examine reprogramming of transcription in response to two different types of cell stress, i.e. oxidative stress and heat shock, we tracked transcription at a nucleotide resolution in mouse embryonic fibroblasts (MEFs) utilizing PRO-seq. Transcribed regulatory regions were identified using the divergent pattern of transcription that characterizes active promoters and enhancers in mammals (12, 13). Enhancers were distinguished from promoters by requiring them to reside over 1 kb from any transcription start site (TSS) of annotated genes. As previously reported (6, 13), the active enhancers identified from PRO-seq profiles, contained enhancer-associated histone marks H3K27ac and H3K4me1 (22, 23) (Fig. S1). To induce oxidative stress, we treated MEFs with different concentrations of menadione, a commonly used ROS generator (24). From the concentrations tested, 30 μM was selected for transcriptional analyses, since it was the lowest concentration that caused oxidative stress, as measured by the decrease in the ratio of reduced and oxidized glutathione (GSG/GSSG) (Fig. S2A). The heat shock response was induced by exposing MEFs to 42°C for 1 h.

Both menadione and heat shock caused remarkable changes in transcription of genes and enhancers (Fig 1A). Interestingly, the changes in transcription were more prominent upon oxidative stress than upon heat shock (Fig. 1A). During both stresses, the number of downregulated genes was greater than the number of upregulated genes, whereas enhancers displayed an opposite pattern (Fig.1A). These results show a general reduction of gene transcription in response to stress, accompanied with increased residency of engaged Pol II at enhancers. Comparison of transcriptional changes at individual genes and enhancers, however, revealed a prominent stress-specific reprogramming of transcription (Fig. 1B).

**Figure 1.**
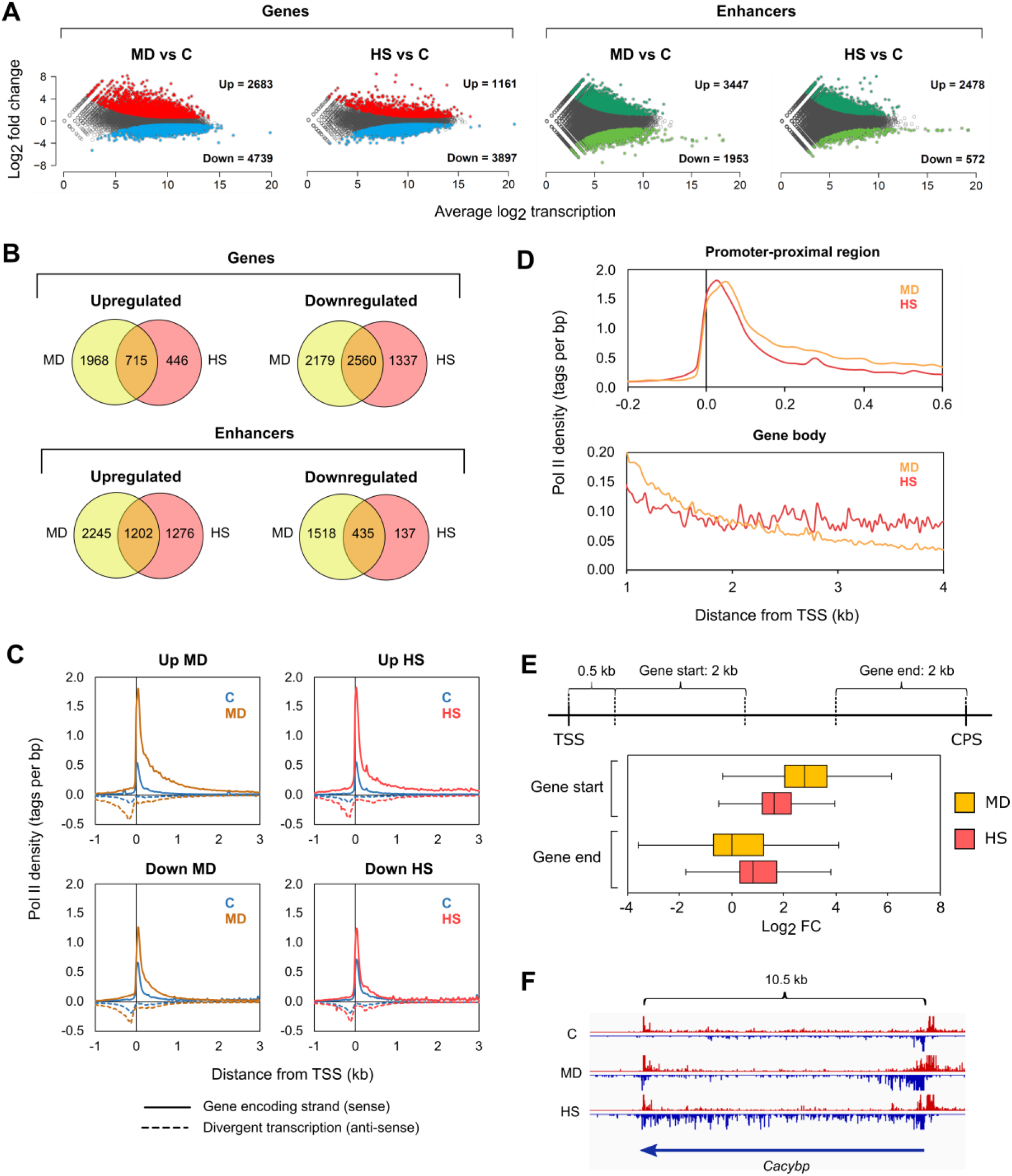
Oxidative stress and heat shock display distinct changes in the transcription of genes and enhancers. (**A**) PRO-seq was performed in MEFs that were exposed to oxidative stress induced by menadione (MD, 30 μM, 2 h) or to heat shock (HS, 42°C, 1 h). The number of upregulated and downregulated genes and enhancers in stressed cells was determined. Threshold for fold change was set to 1.5 and 0.667 to call statistically significant upregulations and downregulations, respectively. (**B**) Genes and enhancers with altered expression during menadione and heat shock were compared to determine the number of genes and enhancers that were upregulated or downregulated in a stress type-specific manner. (**C**) Average density of Pol II was analyzed upstream and downstream of the TSS in the genes that were upregulated or downregulated by menadione or heat shock. Pol II density was measured separately for the sense (solid line) and antisense (dotted line) strands. (**D**) Pol II densities of upregulated genes in menadione and heat shock samples were overlaid in promoter-proximal region (−0.2-0.6 kb relative to the TSS) and gene body (1-4 kb relative to the TSS). (**E**) Log_2_ fold changes (FC) of upregulated genes in cells treated with menadione or heat shock were determined in start and end of the genes. Start of the gene was defined as a 2-kb window starting 0.5 kb downstream from the TSS. End of the gene was defined as a 2-kb window upstream of the CPS. (**F**) PRO-seq profile of calcylin-binding protein (*Cacybp*) gene in cells exposed to menadione and heat shock. C: control.

### Pol II pause-release triggers rapid gene activation in the oxidative stress response

To gain a mechanistic understanding of transcriptional reprogramming, caused by oxidative stress and heat shock, we analyzed the distribution of Pol II along genes and enhancers. Previous studies have shown that upon induction of genes by heat shock, the paused Pol II is released from promoter-proximal regions into elongation simultaneously with the recruitment of new Pol II molecules to the promoters (6, 8, 25). In contrast, repression of gene transcription by heat shock occurs by reducing the pause-release, which causes accumulation of Pol II within promoter-proximal regions (6). Our results show that the distribution of Pol II in the upregulated and downregulated genes follows the same pattern at the promoter-proximal pause region upon menadione treatment and heat shock, indicating that the induction and repression of transcription is regulated at the level of Pol II pause-release during both types of stress (Fig. 1C). These results demonstrate that cells activate and repress stress-specific sets of genes through universal mechanisms.

### Engaged Pol II accumulates at enhancers upon oxidative stress and heat shock

The enhancers that were induced upon stress, showed an absence of Pol II under normal growth conditions (Fig. S2B). Consequently, the critical step in the upregulation of enhancers, upon both oxidative stress and heat shock, was the recruitment of Pol II, which is different from the stress-mediated activation of genes (Fig. S2B). Downregulated enhancers, in turn, displayed Pol II occupancy already under normal growth conditions, and the occupancy decreased in response to both stresses (Fig. S2B). Intriguingly, the profiles of downregulated enhancers, showed several Pol II peaks, which implies that transcriptionally active enhancer clusters, also known as super-enhancers (26), lose engaged Pol II under stress conditions.

### Increased Pol II density at early gene bodies coincides with oxidative DNA damage

A detailed analysis of Pol II distribution along genes revealed that oxidative stress induced a more profound increase in Pol II density at the promoter-proximal region and beginning of the gene body (0-2 kb from TSS) than was detected at heat-activated genes (Fig. 1D). In contrast, as Pol II approached the cleavage and polyadenylation site (CPS) at the 3’-end of the gene, a higher Pol II density was detected in heat-shocked cells (Fig. 1D). Since productive elongation requires Pol II to transcribe through the entire gene body and beyond the CPS, these results suggest a transcriptional hindrance after the release of paused Pol II in the menadione-treated cells. To investigate whether Pol II proceeded to the end of menadione-activated genes, we determined the fold change of engaged Pol II at the start of the gene (0.5–2.5 kb relative to TSS) and the end of the gene (−2–0 kb relative to CPS) (Fig. 1E). We selected the 0.5–2.5 kb region to represent the start of the gene to avoid the paused Pol II from interfering with the measurement of the fold change in the gene body. We also discarded short genes (0–5 kb) from the analysis. Interestingly, menadione caused a greater fold change in the start of the genes than heat shock, while the fold change in the end of the genes was higher upon heat shock (Fig. 1E). These results are exemplified by the calcylin-binding protein (*Cacybp*) gene, which is upregulated by both stresses, but shows elevated levels of Pol II throughout the gene body only upon heat shock (Fig. 1F).

Although the average induction during menadione treatment was observed particularly in the start of the genes, we found that 37% of the menadione-inducible genes included in the analysis, displayed a fold change above 1.5 also in the end of the genes (Fig. S2C). Genes that showed increased levels of Pol II throughout the gene body in menadione-treated cells include fork head box O4 (*Foxo4*) and heme oxygenase 1 (*Hmox1*) (Fig. S2D), known to be critical in the oxidative stress response (27, 28). The induction that was observed only in the start of several menadione-inducible genes could occur due to oxidative DNA damage, which has been shown to impede the elongation of Pol II (29). This is supported by our finding, which shows that the amount of DNA damage, as measured by levels of phosphorylated H2AX, was increased in response to menadione but not heat shock (Fig. S3). Furthermore, the DNA damage is likely to affect open regions, such as early gene bodies where histone acetylation increases upon transcriptional activation (6, 30).

### HSF1 and HSF2 direct the oxidative stress response

HSF1 is a well-known *trans*-activator of protein folding machinery under proteotoxic stress conditions, while the role of HSF2 in the regulation of stress-inducible transcription has remained elusive (17). For determining the specific roles of HSF1 and HSF2 in transcriptional activation of enhancers and genes during oxidative stress and heat shock, we performed PRO-seq in HSF1 knock-out (KO) MEFs and HSF2 KO MEFs, in addition to wild-type (WT) MEFs (Fig. S4A and S4B). To analyze the impact of HSFs on the enhancer transcription, we selected only enhancers, which produced eRNAs and contained either H3K27ac or H3K4me1 mark (Fig. S1). Similar to heat shock, menadione treatment resulted in upregulation of hundreds of genes and enhancers in an HSF1- and/or HSF2-dependent manner (Fig. 2A and 2B). We also found that the transcriptional program was altered in HSF1 and HSF2 KO MEFs already under normal growth conditions (Fig. 2C). This result is in line with the various roles of HSF1 and HSF2 under physiological conditions, including differentiation, development, and cell cycle control as well as in pathological states, such as cancer and neurodegeneration (14, 15).

**Figure 2.**
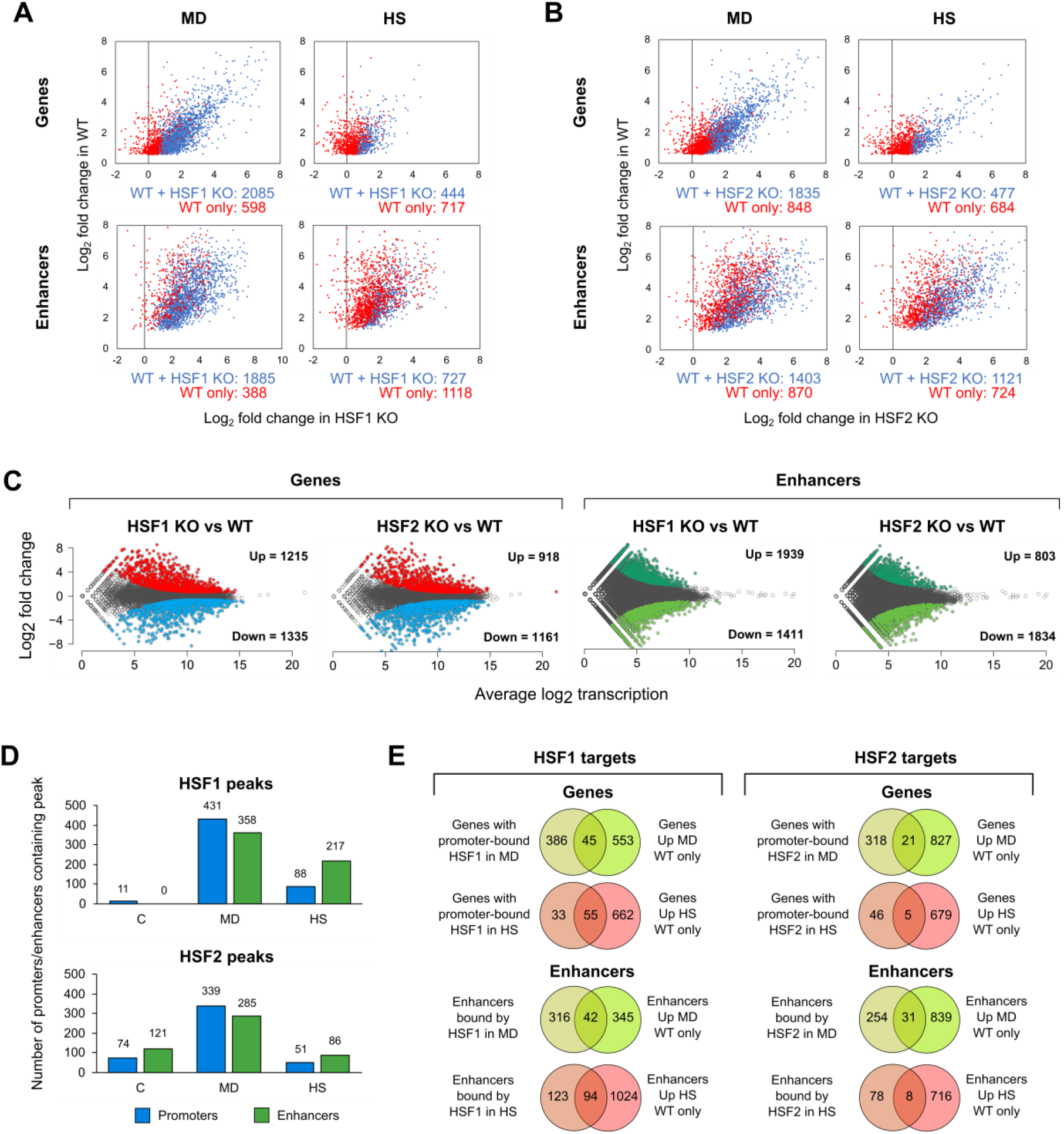
HSF1 and HSF2 reprogram the transcription of genes and enhancers in response to oxidative stress and heat shock. (**A, B**) PRO-seq was performed in wild-type (WT), HSF1 knock-out (HSF1 KO) and HSF2 knock-out (HSF2 KO) MEFs that were exposed to oxidative stress induced by menadione (MD, 30 μM, 2 h) or to heat shock (HS, 42°C, 1 h). Log_2_ fold changes are shown for the genes and enhancers that are upregulated either in WT and KO cells (blue dots) or only WT cells (red dots). Some of the HSF-dependent genes and enhancers are likely false positives, since they displayed high fold change in both WT and KO cells (red dots towards the right side of the panels). In these cases, the fold changes in KO cells were not statistically significant and, therefore, these genes and enhancer are upregulated only in WT cells. (**C**) Comparison between KO and WT cells revealed several genes and enhancers that are upregulated or downregulated in HSF1 and HSF2 KO cells under normal growth conditions. (**D**) Antibodies against HSF1 and HSF2 were used to perform ChIP-seq in MEFs that were exposed to menadione or heat shock. The number of promoters and enhancers that contained HSF1 or HSF2 peak was determined in cells exposed to menadione or heat shock. (**E**) Target genes and enhancers regulated through direct binding of HSF1 or HSF2 were identified by comparing the targets bound by HSF1 or HSF2 with the targets that were upregulated only in WT cells. C: control.

To distinguish the direct targets of HSF1 and HSF2 from the indirect ones, we identified genes and enhancers occupied by HSF1 and HSF2 in stressed cells. We treated WT MEFs with menadione or heat shock, and immunoprecipitated HSF1 and HSF2 for the ChIP-seq analysis. A strong stress-inducible binding of HSF1 to promoters and enhancers was evident during both stresses, and remarkably, the number of HSF1-bound promoters and enhancers was even higher upon menadione treatment than heat shock (Fig. 2D). In addition to HSF1, HSF2 displayed a prominent inducible binding to both promoters and enhancers in menadione-treated cells (Fig. 2D). Unlike HSF1, HSF2 bound to several targets prior to stress exposures, and the number of HSF2 targets did not increase in response to heat shock (Fig. 2D). This observation could be explained by heat-induced degradation of HSF2, which occurs shortly after exposure to heat shock (31). Together, our results indicate distinct kinetics of HSF2-mediated transcription in heat-shocked and ROS-challenged cells.

Next, we identified the direct targets of HSFs whose stress-inducibility was dependent on the binding of HSF1 or HSF2 to the corresponding cis-acting elements in the genome. Our analysis revealed a multitude of menadione- and heat-inducible genes and enhancers, which were dependent on HSF1 binding (Fig. 2E and Table S1). Although menadione-inducbile target genes of HSF1 play roles in various biological processes, many of them were related to protein folding (Table S1). In line with our previous findings (32), HSF2 displayed only few targets during heat shock and was not required for stress-inducible upregulation of HSP genes (Fig. 2E and Table S1). Surprisingly, during oxidative stress, HSF2 was required for the induction of multiple enhancers and genes that are known to regulate protein folding, DNA repair, apoptosis, and other stress-related processes (Fig. 2E and Table S1). This result demonstrates that HSF2 can function, besides a modulator of transcription, also as a *bona fide* transcription factor.

### HSF2 regulates the oxidative stress response independently of HSF1

Since HSF2 has been primarily described as a modulator of HSF1 activity, we sought to understand whether HSF2 can activate transcription independently of HSF1. In menadione-treated cells, we found nine genes and 16 enhancers that were HSF2-specific direct targets (Fig. 3A). One of them is solute carrier family 35 member A5 (Slc35a5) gene, which was induced through binding of HSF2, but not HSF1, to its promoter (Fig. 3C). Thus, we conclude that HSF2 can function as a transcription factor independently of HSF1. Several other menadione-inducible genes and enhancers were regulated by HSF1 and HSF2, or by HSF1 alone, as exemplified by an HSF1-specific target gene, solute carrier family 25 member 38 (*Slc25a38*) (Fig. 3A and 3C). In heat-shocked cells, nearly all HSF targets, including ST13 hsp70 interacting protein (ST13) gene, were regulated by HSF1 alone, although they were occupied by both HSFs (Fig. 3B and 3C). Intriguingly, we identified only five and eight HSF2 target genes and enhancers, respectively, that were regulated also by HSF1 (Fig. 3B). Together, these results show that HSF2 functions as a transcription factor independently of HSF1 during oxidative stress.

**Figure 3.**
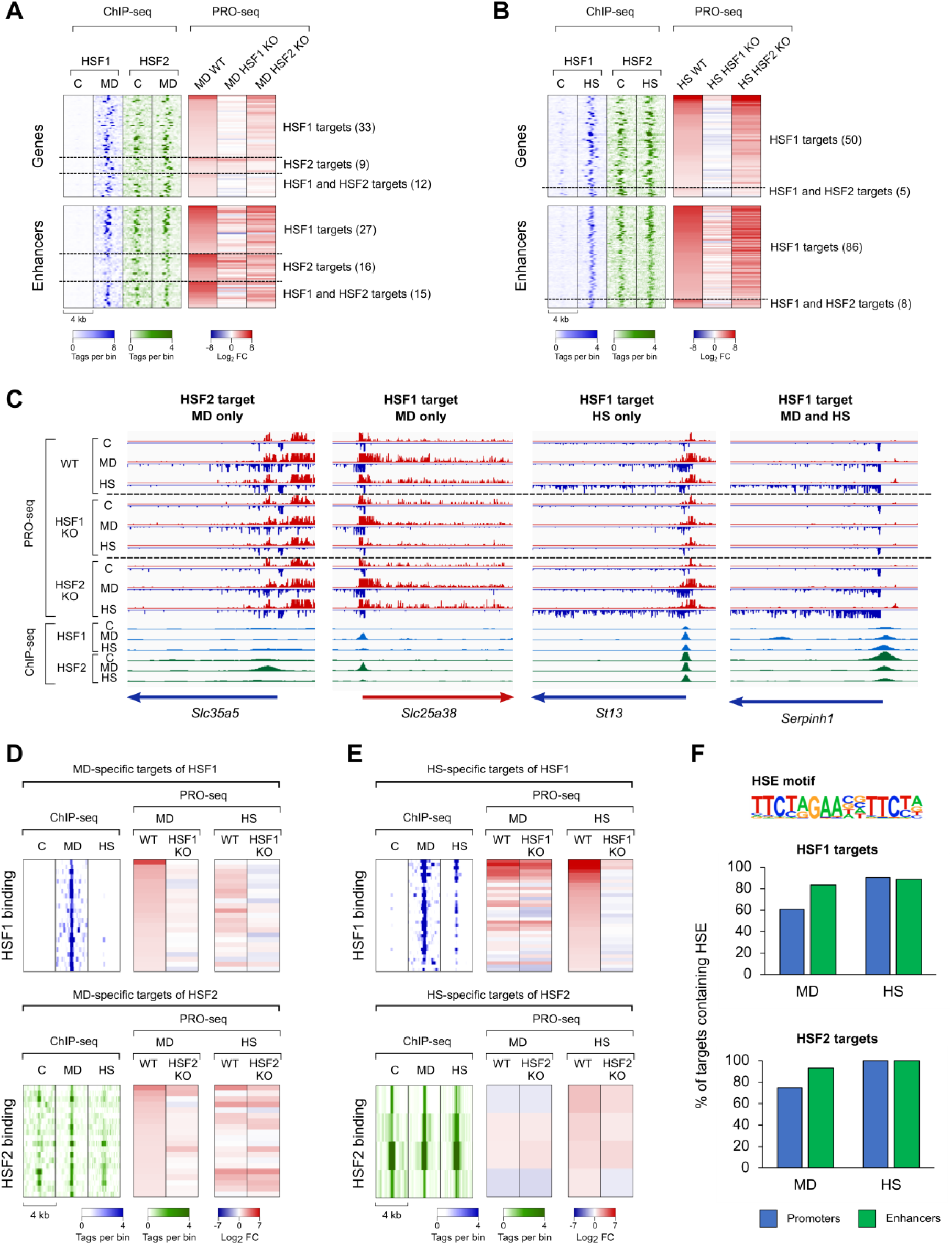
HSF1 and HSF2 drive distinct transcriptional programs upon oxidative stress and heat shock. (**A, B**) Heatmaps were generated from menadione- (MD, 30 μM, 2 h) (A) and heat-treated (HS, 42°C, 1 h) (B) MEFs to show genes and enhancers, which are regulated through direct binding of both HSF1 and HSF2 or only one of these factors. (**C**) PRO-seq and ChIP-seq profiles are shown for selected genes that are induced by HSF1 and HSF2 in response to menadione or heat shock. Headings above each of the four panels indicate whether the gene is regulated by HSF1 or HSF2 during menadione, heat shock or both. (**D, E**) Heatmaps were generated from menadione- (D) and heat shock (E) -specific target genes of HSF1 and HSF2. (**F**) Motif analysis was performed to determine the percentage of menadione- and heat shock-specific targets of HSF1 and HSF2 that contain canonical HSEs. C: control, *Slc35a5:* solute carrier family 35 member A5, *Slc25a38:* solute carrier family 25 member 38, *ST13:* Hsp70 interacting protein, *Serpinh1:* serpin family H member 1.

### HSFs activate distinct transcription programs through stress-specific binding to chromatin

We found that both HSF1 and HSF2 regulated unique sets of genes and enhancers in cells treated with menadione or heat shock (Fig. 3C and Fig. S5). Next, we asked whether HSF1 and HSF2 bind to stress-specific sites in the chromatin to regulate their stress-specific targets. Our results revealed a large group of genes that were occupied and activated by HSF1 and/or HSF2 only in menadione-treated cells, demonstrating for the first time that HSFs can bind unique sites in response to distinct stress stimuli (Fig. 3D). Interestingly, we found that while heat-inducible HSF targets were bound by HSF1 and/or HSF2 also in response to menadione, a majority of these targets were induced in an HSF-dependent manner only in heat-shocked cells (Fig. 3E). This implies that HSFs lack the full *trans*-activation capacity at certain genes during oxidative stress, which could occur either because oxidative stress represses HSFs or because transcriptional co-activators of HSFs are not available during oxidative stress.

Differential binding patterns of HSFs between menadione treatment and heat shock could be explained by their preference for distinct target motifs in the DNA. It is known that HSFs bind to their cis-acting heat shock elements (HSEs), which were originally defined to contain three inverted nGAAn sequences (33). These motifs are called canonical HSEs, but subsequent studies have identified also non-canonical HSEs, which consist of highly variable sequences (34, 35). Therefore, it is plausible that oxidative stress-specific target genes of HSFs contain primarily non-canonical HSEs that are not recognized by current motif finding algorithms. We found that canonical HSEs were less prevalent in the menadione-specific target genes of HSF1 and HSF2 than in their targets specific for heat shock (Fig. 3F). Unlike genes, enhancer targets of both HSFs displayed no differences in the prevalence of canonical HSEs (Fig. 3F). Taken together, our data indicate that HSF1 and HSF2 can bind to cis-acting elements that lack canonical HSEs in their menadione-specific target genes.

### HSF1 and HSF2 bind enhancers to drive stress-inducible gene transcription

Since a majority of HSF1- and HSF2-dependent genes were not directly regulated by promoter-bound HSF1 or HSF2 (Fig. 3E), we hypothesized that these genes could be induced through enhancers. Interestingly, we observed that during heat shock, a prominent number of HSF1-dependent genes resided within 100 kb from the direct enhancer targets of HSF1 (Fig. 4A). Furthermore, most of these genes were devoid of promoter-bound HSF1, suggesting that HSF1 regulates a subset of heat-inducible genes through nearby enhancers (Fig. 4A). Also, several menadione-induced genes required HSFs for activation and had the closest HSF binding-site at a nearby enhancer (Fig. 4A and 4B). However, no general correlation was found between the distance of HSF-dependent genes and the enhancers activated in an HSF-dependent manner upon menadione treatment (Fig. 4A). The importance of HSF2 for stress-induced *trans*-activation at genes and enhancers is illustrated at the beta-1,4-galactosyltransferase 1 (*B4galt1*) locus (Fig. 4B). HSF2 bound stress-inducibly to a distal enhancer within the gene body of B4galt1, and the presence of HSF2 was required both for transcriptional activation at the enhancer and productive elongation at the connected gene (Fig. 4B).

**Figure 4.**
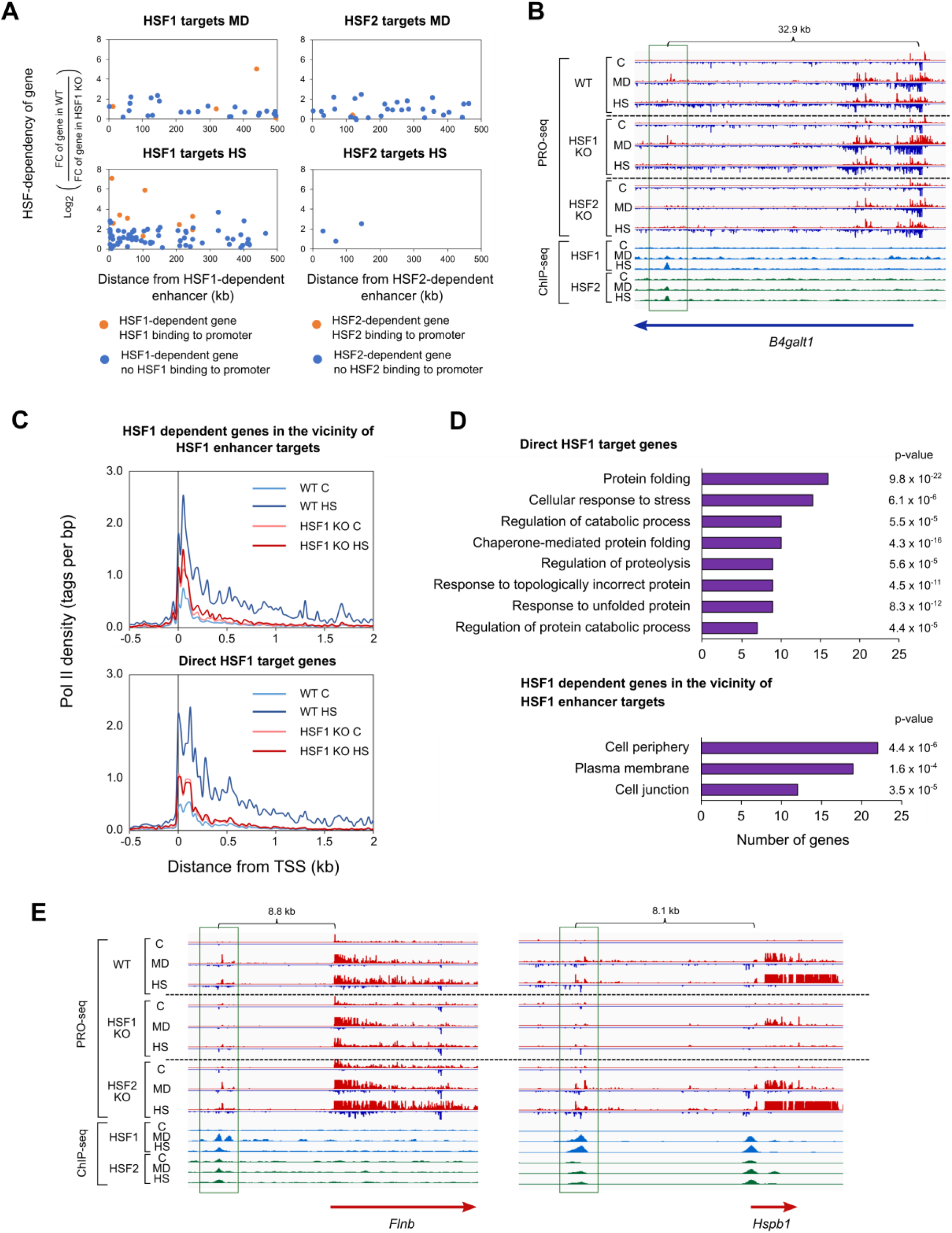
HSF-dependent genes are located in the vicinity of HSF-bound enhancers. (**A**) Distances from the target enhancers of HSF1 and HSF2 to the HSF1- and HSF2-dependent genes were measured in cells exposed to menadione (MD, 30 μM, 2 h) and heat shock (HS, 42°C, 1 h). Distances were calculated between the summit of an enhancer and the TSS of a gene. Genes were divided into two groups depending on whether their promoters were bound by HSF1 or HSF2 (orange dots) or not (blue dots). (**B**) PRO-seq and ChIP-seq profiles of *B4galt1* gene and its downstream enhancer. Enhancer is framed with green rectangle. Enhancer is regulated by direct binding of HSF2, while *B4galt1* gene is devoid of promoter-bound HSF2. (**C**) Average Pol II density was analyzed in the direct HSF1 target genes and HSF1 dependent genes located within 100 kb of direct enhancer targets of HSF1. Pol II densities are shown in wild-type (WT) and HSF1 knock-out (HSF1 KO) MEFs. (**D**) GO terms of two different heat-inducible gene groups were analyzed: direct HSF1 targets and indirect HSF1 targets that were located within 100 kb of direct enhancer targets of HSF1. GO terms were ranked in descending order based on the number of genes associated with each term. (**E**) PRO-seq and ChIP-seq profiles of selected target enhancers and genes of HSF1 that were found in the vicinity of each other. Enhancers are framed with green rectangles. All the enhancers and *Hspb1* gene are regulated through direct binding of HSF1, while *Flnb* gene is devoid of promoter-bound HSF1. C: control, *Flnb:* filamin b, *B4galt1:* beta-1,4-galactosyltransferase 1.

Since only heat-induced target enhancers and genes of HSF1 were found in the vicinity of each other, we assessed how the HSF1-activated enhancers impact distinct steps of transcription at nearby genes during heat shock. Previous studies have shown that binding of HSF1 to promoters is essential for the heat-inducible pause-release and recruitment of Pol II (36, 37). Thus, we analyzed the distribution of Pol II at genes whose heat-induction was indirectly dependent on HSF1 and which were located within 100 kb from direct target enhancers. Our result showed that, similarly to the promoter-bound HSF1, binding of HSF1 to enhancers was required for the pause-release and recruitment of Pol II at nearby genes (Fig. 4C).

Finally, we addressed whether HSF1 regulates different cellular processes through promoters and enhancers in cells exposed to cytotoxic stress, especially heat shock. For this purpose, we compared GO terms between the direct target genes of HSF1 and the indirect target genes located within 100 kb from its enhancer targets. As expected, the direct HSF1 target genes were related to processes of protein folding, proteolysis and cellular stress responses (Fig. 4D). On the contrary, the indirect target genes residing in the vicinity of enhancer targets were strongly associated with GO terms, such as cell periphery, plasma membrane and cell junction (Fig. 4D). Examples of these targets are filamin b (*Flnb*) and membrane-associated guanylate kinase, WW and PDZ domain containing 1 (*Magi1*) genes, both of which encode proteins localized to the plasma membrane (Fig. 4E and Fig. S6). Furthermore, certain genes with the highest transcriptional induction, e.g. Hspb1, recruited HSF1 both to the promoter and a nearby enhancer (Fig. 4E).

Previous studies have shown that besides protein folding, HSFs regulate genes related to many other processes, including cell adhesion (38, 39). Moreover, maintenance of cell adhesions was shown to be essential for surviving stress (39). Our results advance these studies by revealing that in contrast to the promoter-bound HSF1, which drives the classical chaperone genes, binding of HSF1 to enhancers activates genes encoding proteins localized at cell junctions and the plasma membrane. We also found that both HSF1 and HSF2 are important for the activation of oxidative stress-inducible genes and enhancers, which are different from heat shock-inducible HSF targets. Hereby, we conclude that HSFs function as multi–stress-responsive transcription factors that activate distinct sets of genes and enhancers depending on the type of stress experienced by cells.

## DISCUSSION

Mechanisms of transcriptional reprogramming in response to cellular stresses, especially acute heat shock, are well characterized, but they have remained poorly understood under other stress conditions. Here, we provide the first comprehensive study, in which we combined PRO-seq and ChIP-seq to determine the roles of HSF1 and HSF2 in the regulation of nascent transcription in cells exposed to two different types of cytotoxic stress, i.e. oxidative stress and heat shock. As illustrated in our model (Fig. 5), these two stresses cause clearly stress type-specific changes to the transcription of genes and enhancers. Although the transcriptional programs differ between oxidative stress and heat shock, our results reveal that during both stresses, genes are regulated at the level of Pol II pause-release, while enhancers are regulated via recruitment of Pol II. Unlike heat-inducible genes, a large fraction of oxidative stress-inducible genes displayed elongating Pol II only within the early gene body (0-2 kb from TSS). This could be due to oxidative DNA damage, which has been shown to cause stalling of elongating Pol II (29). Other possible explanations are a slower movement speed of Pol II and a failure in the chromatin remodeling in front of elongating Pol II during oxidative stress.

**Figure 5.**
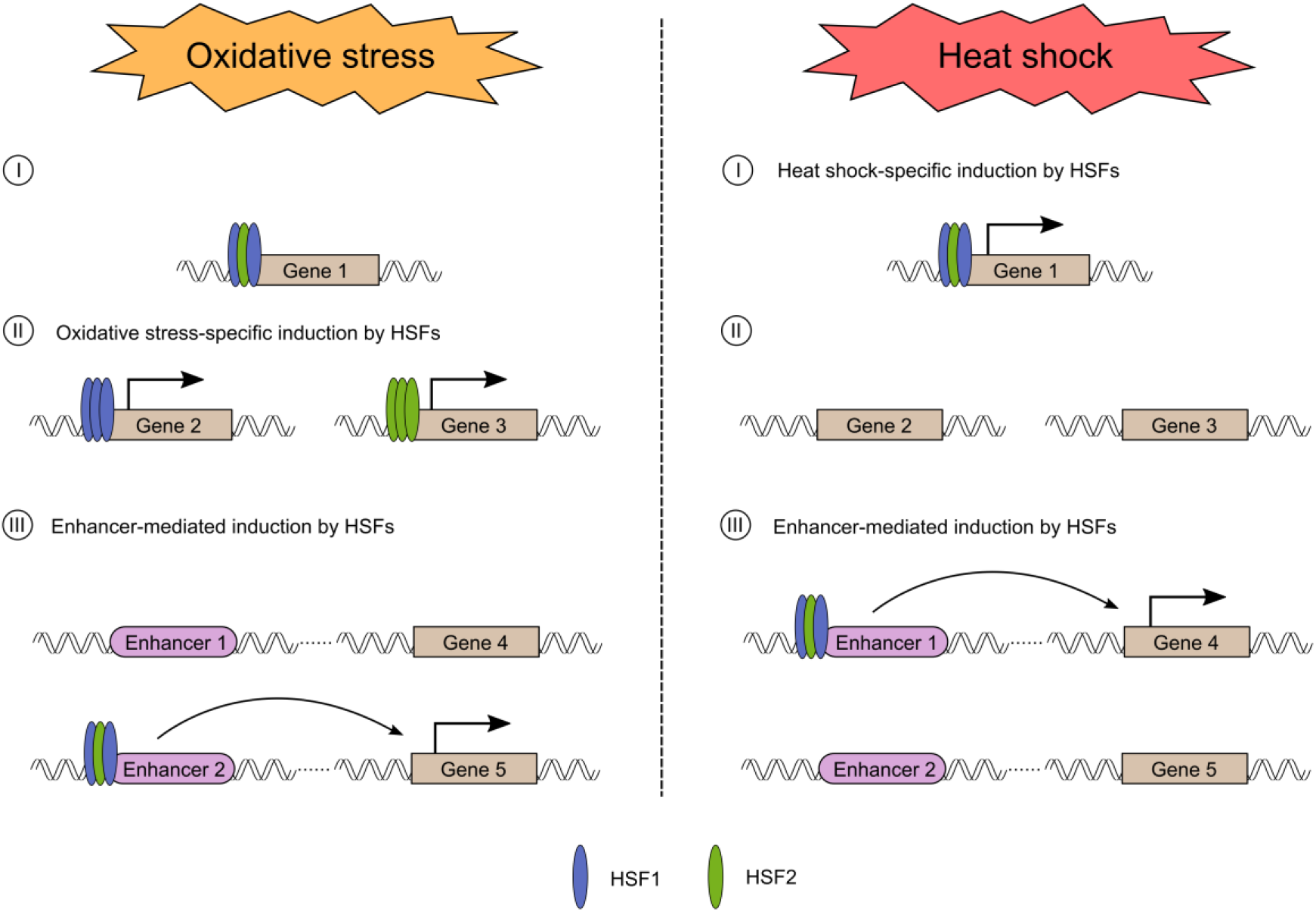
Schematic model of how HSF1 and HSF2 drive stress-specific transcriptional programs through activation of genes and enhancers. (I) HSF1 and HSF2 co-occupy several gene promoters during oxidative stress and heat shock. However, many of these HSF1 and HSF2 - bound genes are only induced in response to heat shock, in an HSF1-dependent manner. (II) Increased levels of ROS trigger HSF1 and HSF2 to bind to their oxidative stress-specific target genes. In contrast to heat shock, oxidative stress induces HSF2 to localize to several gene promoters and activate transcription independently of HSF1. (III) HSF1 and HSF2 bind stress-inducibly to a large number of enhancers. The HSF-bound enhancers differ in heat shock *versus* oxidative stress, but during both conditions HSFs can trigger the release of paused Pol II from promoter-proximal region of a near-by gene. Please note, in this model co-occupancy of HSF1 and HSF2 is drawn as a heterotrimer.

Transcriptional regulation in oxidative stress responses has been largely devoted to nuclear factor erythroid 2-related factor 2 (Nrf2) and members of the Foxo family (27, 40). Here, we expand the repertoire of transcription factors in oxidative stress by identifying HSF1 and HSF2 as new regulators of genes and enhancers in cells exposed to elevated ROS production (Fig. 5). This is an important finding, since (i) HSFs have been considered as master regulators of proteotoxic stress responses, especially the heat shock response, and (ii) HSF2 has been reported to act only as a modulator of HSF1 activity in heat-shocked cells (32). After 30 years of the cloning of the mammalian HSFs (41–43), we now provide the first evidence that HSF2 functions as a *bona fide* transcription factor independently of HSF1, and that HSF2 has a prominent impact on the oxidative stress response. We also show that both HSFs bind and regulate largely different targets upon oxidative stress and heat shock (Fig. 5). Intriguingly, many of the oxidative stress-specific targets genes of HSF1 and HSF2 lacked canonical HSEs, and the mechanism which recruits HSF1 and HSF2 to these atypical sites remains to be established. It is likely that HSFs bind to their oxidative stress-specific targets by interacting with cofactors that are activated by changes in the cellular redox status. Formation of these interactions, in turn, could involve stress-specific protein modifications, since HSFs are known to undergo extensive post-translational modifications, including the oxidation of two redox-sensitive cysteines within the DNA-binding domain of HSF1 (17, 18).

Our data uncover a new regulatory level of stress-inducible transcription that is mediated through enhancers, which in turn are activated by HSFs (Fig. 5). We found that unlike promoter-bound HSF1, which activates classical chaperone genes, enhancer-bound HSF1 was required for the transcriptional induction of cell type-specific genes, including genes that encode proteins localized in the plasma membrane and cell junctions. Enhancer-mediated induction of genes by HSFs is likely not restricted to stress, since HSFs are important transcription factors in a wide variety of physiological processes, including development, differentiation, and metabolism, as well as pathologies, especially cancer and neurodegeneration (14, 15). Furthermore, enhancers play key roles in determining cell fate during development and differentiation, while cancer cells hijack oncogenic enhancers to promote malignancy (44). In future studies, it will be fundamental to determine the functional relevance of HSF-activated enhancers in physiology and pathology.

## DATA AVAILABILITY

Collection of PRO-seq and ChIP-seq raw data has been deposited to Gene Expression Omnibus (GEO) database with accession number GSE183245. In addition to raw data, accession contains bedgraph files that are used for the visualization of the data.

## ACKNOWLEDGEMENT

We thank all the members of Sistonen laboratory for constructive discussions during the preparation of the manuscript. The Finnish Functional Genomics Centre (Turku Bioscience, University of Turku, Åbo Akademi University and Biocenter Finland) is acknowledged for services, instrumentation, and expertise.

