## Supplementary data for "HSFs drive stress type-specific transcription of genes and enhancers"

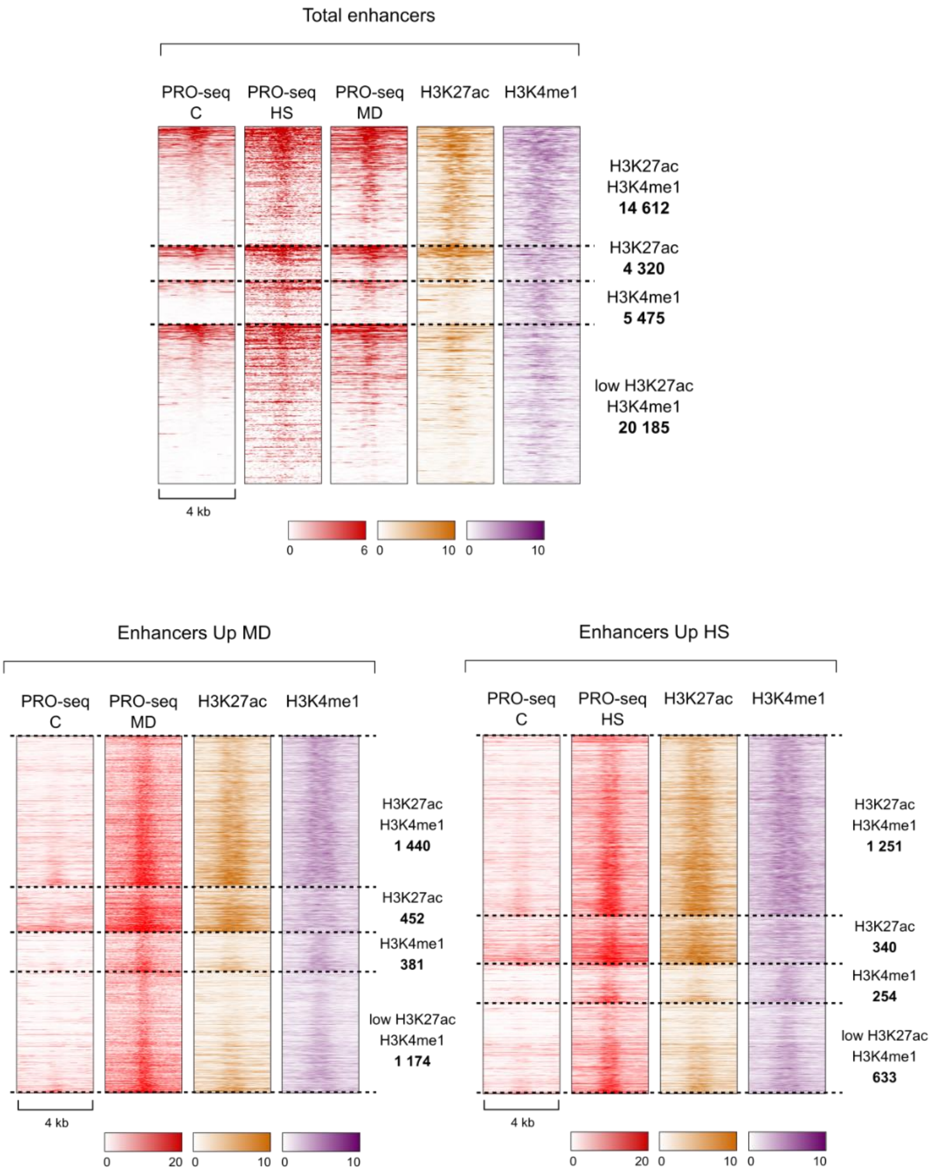


**Figure S1. Enhancers detected by PRO-seq contain enhancer-associated histone marks, H3K27ac and H3K4me1.** PRO-seq was used to identify transcriptionally active enhancers in MEFs treated with menadione (MD, 30 µM, 2 h) or heat shock (HS, 42°C, 1 h). Enhancers were analyzed for their content of histone marks H3K27ac and H3K4me1, both of which are known to be enriched in enhancers (22, 23). Histone marks were analyzed separately from total enhancers and upregulated enhancers. Intensity of the signal in the heatmaps indicates the number of tags per bin. Bin size was set to 50 bp. C: control.

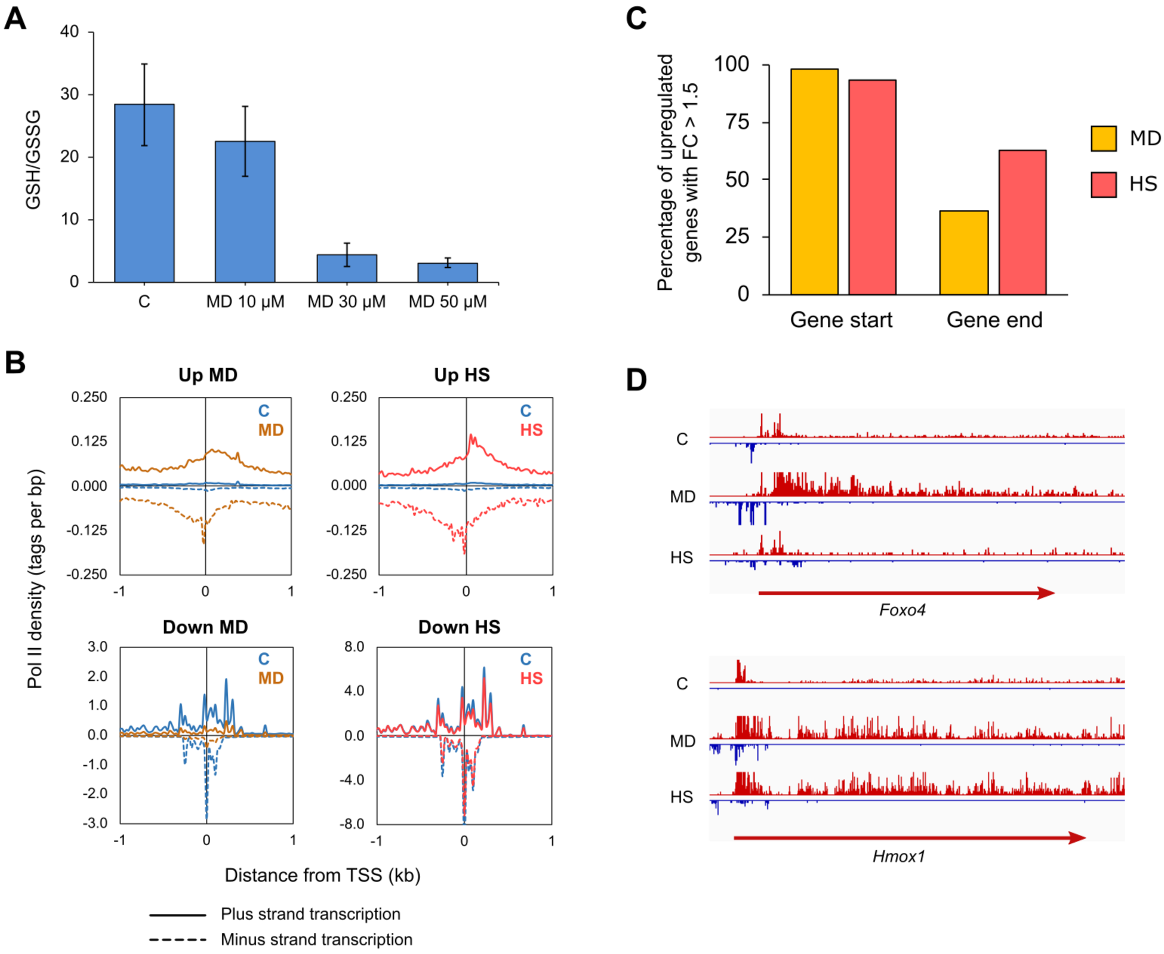


**Figure S2. Profiles of Pol II are similar in enhancers but different in genes between oxidative stress- and heat-treated cells.** (**A**) MEFs were treated with different concentration of menadione for 2 hours to induce oxidative stress. Next, the level of oxidative stress was assessed by measuring the ratio between reduced and oxidized glutathione (GSH/GSSG). (**B**) PRO-seq was performed in MEFs that were exposed to oxidative stress induced by menadione (MD, 30 µM, 2 h) or to heat shock (HS, 42°C, 1 h). Average density of Pol II was analyzed within enhancers that were upregulated or downregulated by menadione or heat shock. Pol II density was measured separately for plus (solid line) and minus (dotted line) strands. (**C**) Fold changes (FC) of upregulated genes in menadione- and heat-treated cells were determined in start and end of the genes. After this, the percentage of genes that displayed FC over 1.5 in gene start or end were calculated. Start of the gene was defined as a 2 kb window starting 0.5 kb downstream from TSS. End of the gene was defined as a 2kb window upstream of TTS. (**D**) PRO-seq profiles of *Foxo4* and *Hmox1* genes in cells exposed to menadione and heat shock. C: control, *Foxo4*: fork head box O4, *Hmox1*: and heme oxygenase 1.

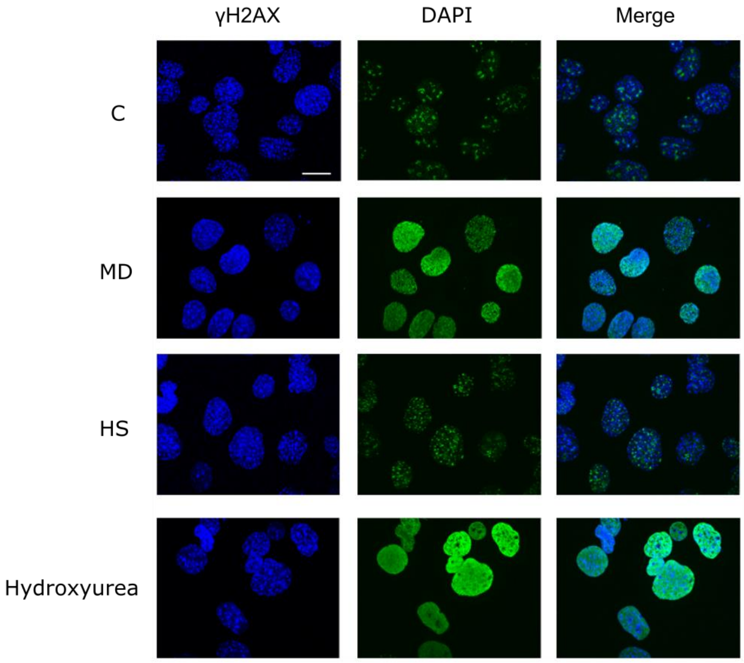


**Figure S3. Oxidative stress induced by menadione causes DNA damage.** MEFs were exposed to oxidative stress induced by menadione (MD, 30 µM, 2 h) or to heat shock (HS, 42°C, 1 h). The amount of DNA damage was determined by immunofluorescence staining of the phosphorylated H2AX (γH2AX). Hydroxyurea (2 mM, 17 h) was used as a positive control to induce DNA damage. DAPI was used to stain DNA. All images correspond to maximum intensity projections. Scalebar: 20 µm. C: control.

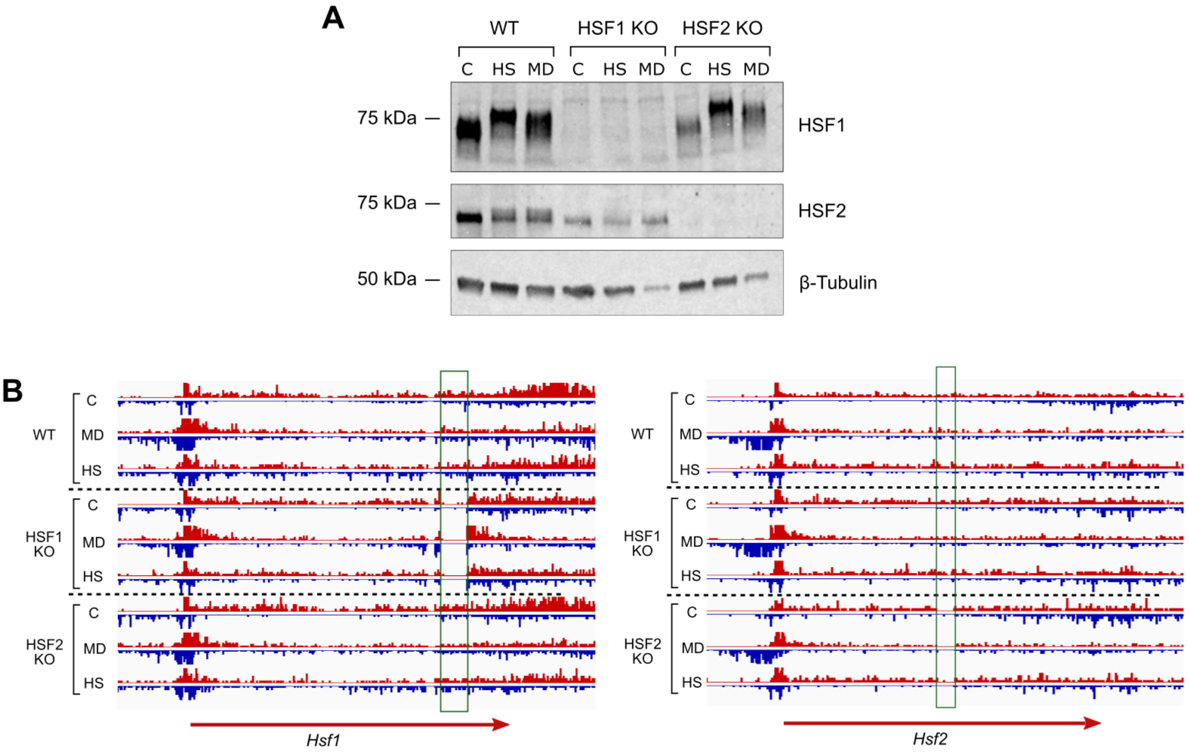


**Figure S4. Validation of HSF1 and HSF2 knock out MEFs.** (**A**) Western blot was used to determine the levels of HSF1 and HSF2 in wild-type (WT), HSF1 knock-out (HSF1 KO) and HSF2 knock-out (HSF2 KO) MEFs that were exposed to menadione (MD, 30 µM, 2 h) or to heat shock (HS, 42°C, 1 h). β-tubulin was used as a loading control. (**B**) PRO-seq profiles of *Hsf1* and *Hsf2* genes in WT, HSF1 KO and HSF2 KO MEFs that were exposed to menadione or heat shock. Green rectangles indicate regions of HSF1 and HSF2 genes that were deleted to create KO mice, from which the cell lines used in this study have been derived. C: control.

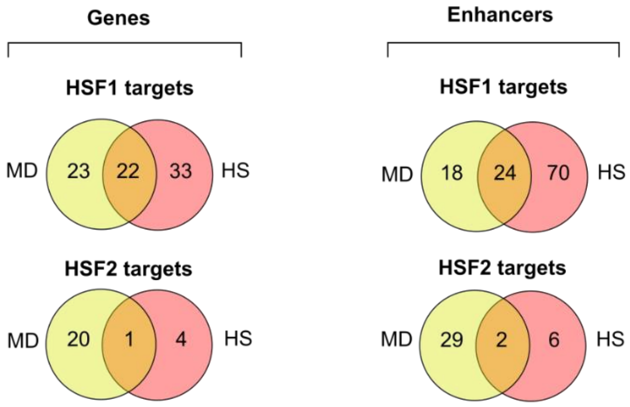


**Figure S5. HSF1 and HSF2 regulate stress-specific sets of genes and enhancers.** Comparison between menadione- and heat-inducible targets of HSF1 and HSF2 revealed genes and enhancers that are regulated by HSF1 and HSF2 in a stress type-specific manner.

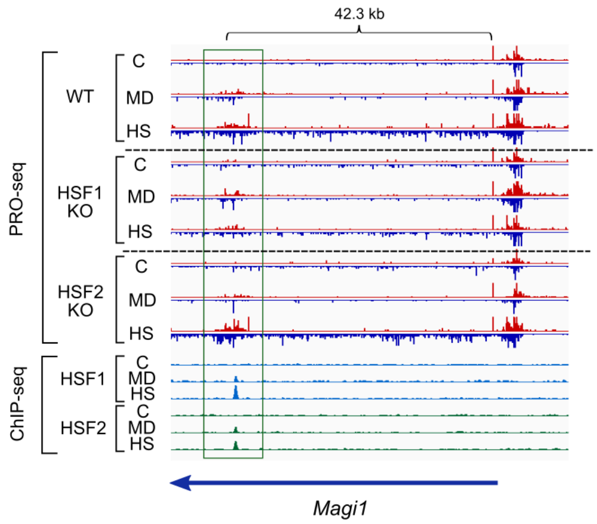


**Figure S6. Direct target enhancer of HSF1 is located in the vicinity of the HSF1-dependent gene, *Magi1.*** PRO-seq and ChIP-seq profiles of *Magi1* gene and its downstream enhancer. Enhancer is framed with green rectangle. Enhancer is regulated by direct binding of HSF1, while *Magi1* gene is devoid of promoter-bound HSF1. C: control, *Magi1*: membrane associated guanylate kinase, WW and PDZ domain containing 1; MD: menadione, 30 µM, 2 h; HS: heat shock, 42°C, 1 h.

| HSF1 target genes in HS | | HSF1 target genes in MD | | HSF2 target genes in HS | | HSF2 target genes in MD | |
|---|---|---|---|---|---|---|---|
| Gene name | FC in WT | Gene name | FC in WT | Gene name | FC in WT | Gene name | FC in WT |
| Hspa1a | 333.5 | Hspa1l | 42.6 | Adgra3 | 3.2 | Phlda1 | 9.6 |
| Hspb1 | 287.5 | Mest | 32.8 | Ints2 | 2.2 | Cetn4 | 4.3 |
| Hspa1b | 120.7 | Lrrc61 | 6.7 | Xpnpep3 | 1.9 | Bst2 | 3.4 |
| Dnaja4 | 52.4 | Hikeshi | 4.8 | Abcc5 | 1.9 | Gcnt2 | 2.7 |
| Hspa1l | 25.6 | Rbm42 | 4.7 | Acot7 | 1.5 | Gm4285 | 2.5 |
| Dnajb1 | 22.4 | Mns1 | 4.4 | | | Plin2 | 2.4 |
| Hspe1 | 19.8 | Slc25a38 | 4.3 | | | Scamp3 | 2.2 |
| Hspa4l | 15.9 | Nfkbid | 4.1 | | | Abcc5 | 2.0 |
| Bag3 | 14.5 | Bst2 | 3.4 | | | Actr5 | 1.9 |
| Mns1 | 14.1 | Zscan29 | 3.3 | | | Slc35a5 | 1.9 |
| Swt1 | 9.5 | Gm10069 | 3.3 | | | AW549877 | 1.8 |
| Esr1 | 7.2 | Gm13830 | 3.2 | | | Chrac1 | 1.8 |
| Hspd1 | 7.1 | Kctd18 | 3.1 | | | Txnip | 1.8 |
| Serpinh1 | 6.8 | Saraf | 3.0 | | | Ppid | 1.8 |
| Gm10069 | 6.7 | Rras | 2.9 | | | Mcoln1 | 1.7 |
| Hspb8 | 6.2 | Cacybp | 2.9 | | | Cnpy4 | 1.7 |
| Stip1 | 6.0 | B4galt2 | 2.7 | | | Serpinh1 | 1.7 |
| Uspl1 | 5.0 | Gcnt2 | 2.7 | | | P4ha1 | 1.6 |
| Hspa8 | 4.8 | Tmem33 | 2.6 | | | Ube2g2 | 1.6 |
| Lman2l | 4.2 | Arl6ip4 | 2.4 | | | Parp3 | 1.6 |
| Ahsa1 | 4.2 | Hspa4l | 2.4 | | | Aptx | 1.6 |
| Kctd18 | 4.2 | Hsp90ab1 | 2.2 | | | | |
| Hsp90ab1 | 4.1 | Stip1 | 2.2 | | | | |
| P4ha1 | 4.1 | Ube2b | 2.2 | | | | |
| Hikeshi | 4.1 | Hspd1 | 2.0 | | | | |
| Cacybp | 4.0 | Chordc1 | 2.0 | | | | |
| Fkbp4 | 4.0 | Dbndd2 | 2.0 | | | | |
| Chordc1 | 3.9 | Abcc5 | 2.0 | | | | |
| Mrfap1 | 3.8 | Atp6v1a | 1.9 | | | | |
| St13 | 3.7 | Actr5 | 1.9 | | | | |
| Tmem33 | 3.7 | Cct7 | 1.9 | | | | |
| Ubqln1 | 3.5 | Slc35e2 | 1.8 | | | | |
| Adgra3 | 3.2 | Chrac1 | 1.8 | | | | |
| Trmt1l | 3.0 | Fkbp4 | 1.8 | | | | |
| Chchd2 | 2.8 | Ppid | 1.8 | | | | |
| Gm6297 | 2.8 | Mcoln1 | 1.7 | | | | |
| Azi2 | 2.6 | Cnpy4 | 1.7 | | | | |
| Slc35e2 | 2.3 | Swt1 | 1.7 | | | | |
| Pradc1 | 2.3 | Serpinh1 | 1.7 | | | | |
| Snx3 | 2.2 | Azi2 | 1.7 | | | | |
| Ints2 | 2.2 | P4ha1 | 1.6 | | | | |
| Rab32 | 2.2 | Zfp46 | 1.6 | | | | |
| Lrrc61 | 2.2 | Ube2g2 | 1.6 | | | | |
| Particl | 2.2 | Vipas39 | 1.6 | | | | |
| Serpinb6a | 2.1 | Aptx | 1.6 | | | | |
| Xpnpep3 | 1.9 | | | | | | |
| Abcc5 | 1.9 | | | | | | |
| Fbxl14 | 1.9 | | | | | | |
| Prss23 | 1.8 | | | | | | |
| Trim65 | 1.8 | | | | | | |
| Vipas39 | 1.8 | | | | | | |
| Ubb | 1.8 | | | | | | |
| Dennd1b | 1.7 | | | | | | |
| Cct7 | 1.6 | | | | | | |
| Acot7 | 1.5 | | | | | | |

**Table S1. List of oxidative stress- and heat shock-inducible target genes of HSF1 and HSF2.** Combination of PRO-seq and ChIP-seq was used to identify genes in MEFs that are regulated through direct binding of HSF1 and HSF2 during oxidative stress induced by menadione (MD, 30 µM, 2 h) or heat shock (HS, 42°C, 1 h). Genes in the list are ranked in the descending order according to their fold change (FC) in wild-type MEFs.
